# Stochastic model of vesicular stomatitis virus replication reveals mutational effects on virion production

**DOI:** 10.1101/2023.07.24.550258

**Authors:** Connor R. King, Casey-Tyler Berezin, Jean Peccoud

**Author notes:** Corresponding author: Jean Peccoud.

## Abstract

We present the first complete stochastic model of vesicular stomatitis virus (VSV) intracellular replication. Previous models developed to capture VSV’s intracellular replication have either been ODE-based or have not represented the complete replicative cycle, limiting our ability to understand the impact of the stochastic nature of early cellular infections on virion production between cells and how these dynamics change in response to mutations. Our model accurately predicts changes in mean virion production in gene-shuffled VSV variants and can capture the distribution of the number of viruses produced. This model has allowed us to enhance our understanding of intercellular variability in virion production, which appears to be influenced by the duration of the early phase of infection, and variation between variants, arising from balancing the time the genome spends in the active state, the speed of incorporating new genomes into virions, and the production of viral components. Being a stochastic model, we can also assess other effects of mutations beyond just the mean number of virions produced, including the probability of aborted infections and the standard deviation of the number of virions produced. Our model provides a biologically interpretable framework for studying the stochastic nature of VSV replication, shedding light on the mechanisms underlying variation in virion production. In the future, this model could enable the design of more complex viral phenotypes when attenuating VSV, moving beyond solely considering the mean number of virions produced.

**Author Summary:** This study presents the first complete stochastic model of vesicular stomatitis virus (VSV) replication. Our model captures the dynamic process of VSV’s replication within host cells, accounting for the stochastic nature of early cellular infections and how these dynamics change in response to mutations. By accurately predicting changes in mean virion production and the distribution of viruses in gene-shuffled VSV variants, our model enhances our understanding of viral replication and the variation we see in virion production. Importantly, our findings shed light on the mechanisms underlying the production of VSV virions, revealing the influence of factors such as the duration of the early infection phase and the interplay between the genome’s ability to switch into an inactive state and viral protein production. We go beyond assessing the mean number of virions produced and examine other effects of mutations, including the probability of aborted infections and the variability in virion production. This stochastic model provides a valuable framework for studying the complex nature of viral replication, contributing to our understanding of single-cell viral dynamics and variability. Ultimately, this knowledge could pave the way for designing more effective strategies to attenuate VSV.

## Introduction

Vesicular stomatitis virus (VSV) is a non-segmented negative-strand RNA virus (NNSV) that has been studied for years as an agricultural pest. It is a model for negative-sense RNA viruses and a biomedical tool (1, 2, 3). VSV’s extensive characterization makes it an ideal system for model-based investigation of specific viral mechanisms of NNSVs and broader questions about viral behaviors. Since 2000, several mechanistic models have been developed to dissect different characteristics of VSV. Recently, a model of transcription in VSV and respiratory syncytial virus (RSV) revealed a novel theory for why the transcription of genes along the genome follows a predictable gradient dependent on position (4). This model showed that the long-held understanding of how NNSV transcription works, assuming a single promoter at the 3’ end of the genome and a constant rate of attenuation between genes (1, 5), could not produce the data that we see in the literature and led to the development of an entirely new model consistent with these observations.

In this study, we seek to understand how the stochastic nature (6) of the reactions that occur during intracellular replication contributes to the variation we see in the number of viruses produced both between different cells and between different variants. Previous models of the dynamics of intracellular replication in VSV have fallen short of capturing the stochastic behavior of virion production while remaining biologically interpretable. The first model was an ordinary differential equation (ODE)-based model made to identify methods to attenuate VSV (7). This model predicted how balancing transcription and genome synthesis rates might affect viral growth and was later built upon by predicting how all 120 possible gene-shuffled variants of VSV might behave (8). The model, while thorough, is extremely complex, making it difficult to interpret and work with. In addition, the model is deterministic and, therefore, unable to capture cell-to-cell variation in virion production.

The next model of VSV intracellular replication sought to simulate the cell-to-cell variation of virion production (9). This model introduces a new hybrid simulation algorithm that includes a new stochastic simulation algorithm called a delayed stochastic simulation algorithm (DSSA). The model of the VSV life cycle has been limited to transcription, translation, and replication reactions; it lacks reactions corresponding to the assembly of the viral particles. In addition, while the model does make a distinction between active and inactive RNA genomes, which is biologically accurate, the active genome is assumed to be the one that is not bound to any proteins. This is inaccurate as the nucleoprotein (N protein) is an essential participant in the facilitation of polymerase activity and it is rare to find genomes that are not encapsidated to some degree (10, 11). Furthermore, the model assumed that the transcription rate of genes is a function of the number of genomes and L polymerase proteins, which was later disproven (12). As a result of these simplifications, the model validation is limited to a qualitative comparison of the dynamics of the variables representing genome numbers with the known dynamics of virus production.

The third model was an ODE model that modeled the production of N protein encoding mRNA molecules as a function of the number of genomes present in the cell. Here it was shown that the number of mRNA molecules is purely a function of the number of genomes and that the L polymerase does not need to be considered. This is consistent with the observation that VSV forms phase-separated compartments (13, 14), and that it is normal for viruses to form factories which may be uninhibited by fluctuations in the concentration of replicase (15, 16). However, for our purposes, this model is too simplistic and does not consider stochasticity. A fourth stochastic model was also developed that only sought to replicate extremely early infection dynamics (less than 20 seconds into the infection) purely to compare parameter estimation methods (17). This model was too limited for our purposes and did not incorporate the creation of new virions as well.

Here we present the first stochastic model of VSV that incorporates the entire replication cycle. The model relies only on mass action reactions, meaning the only factors considered when calculating the reaction rate are the reactant concentrations, and incorporates the synthesis, degradation, and assembly reactions of VSV and can easily be simulated with the standard Gillespie stochastic simulation algorithms. The model also incorporates a transition of the viral genomes from an active to an inactive state facilitated by the M protein (1, 2, 18). The model is the most accurate to date for predicting changes in mean virion production in gene shuffled VSV variants (19) and can additionally capture the distribution of the number of viruses produced by VSV. While we began working with a model that incorporated the individual binding of every single protein that binds to VSV, we ultimately developed a stochastic model that treats the amount of protein that binds to the virus as a single unit. This is consistent with observations that the number of proteins incorporated into a virion varies, such that having an exact number of proteins bind to the genome is not necessarily biologically accurate (20). We also know that some proteins that make up the virion, like the G protein, assemble independently of the other components before being incorporated into the virion (21). We believe this to be the best approach to modeling the dynamics of VSV replication, as we capture the stochastic behavior of virion assembly in this model while producing the most accurate predictions to date for the influence of mutations on virion production. This model allowed us to improve our understanding of both intercellular variation and variation between variants in virion production. Intercellular variability appears to be the result of how long it takes for the virus to exit the early phase of infection. However, the differences between variants instead appear to arise from the complexity of balancing the time the (-) genome spends in the active state and the speed of incorporating new (-) genomes into virions, which in turn is balanced by the production of viral components.

## Results

### Model of VSV Replication

The full model can be found in S2 Dataset under the name Full_Model.xml.

### Molecular species

All species abbreviations are defined in Table 1. A single VSV particle consists of 1 negative-sense strand of RNA genome (nsg), 1258 N, 466 P, 1826 M, 1,205 G, and 50 L (22). When the RNA genome is released into the cell, the M protein travels to the nucleus while the N, P, and L proteins stay associated with the genome as an RNP complex. As such, we set the initial conditions as a single nsg_N molecule with 466 P, 1,205 G, and 50 L available, the quantity of N to zero since it is already bound to the genome, and M to zero as well since it is in the nucleus and not available in the cytoplasm.

**Table 1.** List of abbreviations used for each molecular species in the model.

| Molecular Species | Abbreviation |
| --- | --- |
| Negative-Sense Genome (Free Genome) | nsg |
| Positive-Sense Genome (Free Antigenome) | psg |
| Nucleoprotein | N |
| N bound genome | nsg_N |
| N bound antigenome | psg_N |
| Phospho-protein cofactor | P |
| N and P bound genome | nsg_NP |
| RNA-dependent RNA polymerase | L |
| N, P and L bound genome | nsg_NPL |
| Matrix protein | M |
| N, P, L, and M bound genome | nsg_NPLM |
| Glycoprotein | G |
| N, P, L, M, and G bound genome | nsg_NPLMG |
| Nucleoprotein mRNA | mN |
| Phospho-protein cofactor mRNA | mP |
| Matrix protein mRNA | mM |
| Glycoprotein mRNA | mG |
| RNA-dependent RNA polymerase mRNA | mL |
| Assembled Viral Particle | V |
| Either nsg, nsg_N, nsg_NP, nsg_NPL | Active nRNP |
| Either nsg_NPLM, nsg_NPLMG, and V | Inactive nRNP |
| Either psg or psg_N | pRNP |

### Assembly

In this model, assembly occurs sequentially, starting with the binding of the N proteins to the nsg, followed by subsequent binding of the P proteins, the L proteins, the M proteins, and finally the G proteins (Fig 1). This was inspired by the three layers of the VSV virion with the most internal layer consisting of a single molecule of nsg_NPL which is encased in an outer layer of M proteins and wrapped in a membrane studded with G proteins (1, 18). The binding of each of these factors is broken into several steps, with a single protein binding at each step and each step having the same rate constant listed next to the overall reaction in the table. In this model, psg only recruits the N protein. The literature-derived reaction rates were taken from the binding rate of the N protein from a previous model (9) where the binding rate is assumed to be 0.0461 for the binding of one molecule of N to the genome/antigenome. Here, we make this our initial guess for the binding rate of all proteins.

**Fig 1.**
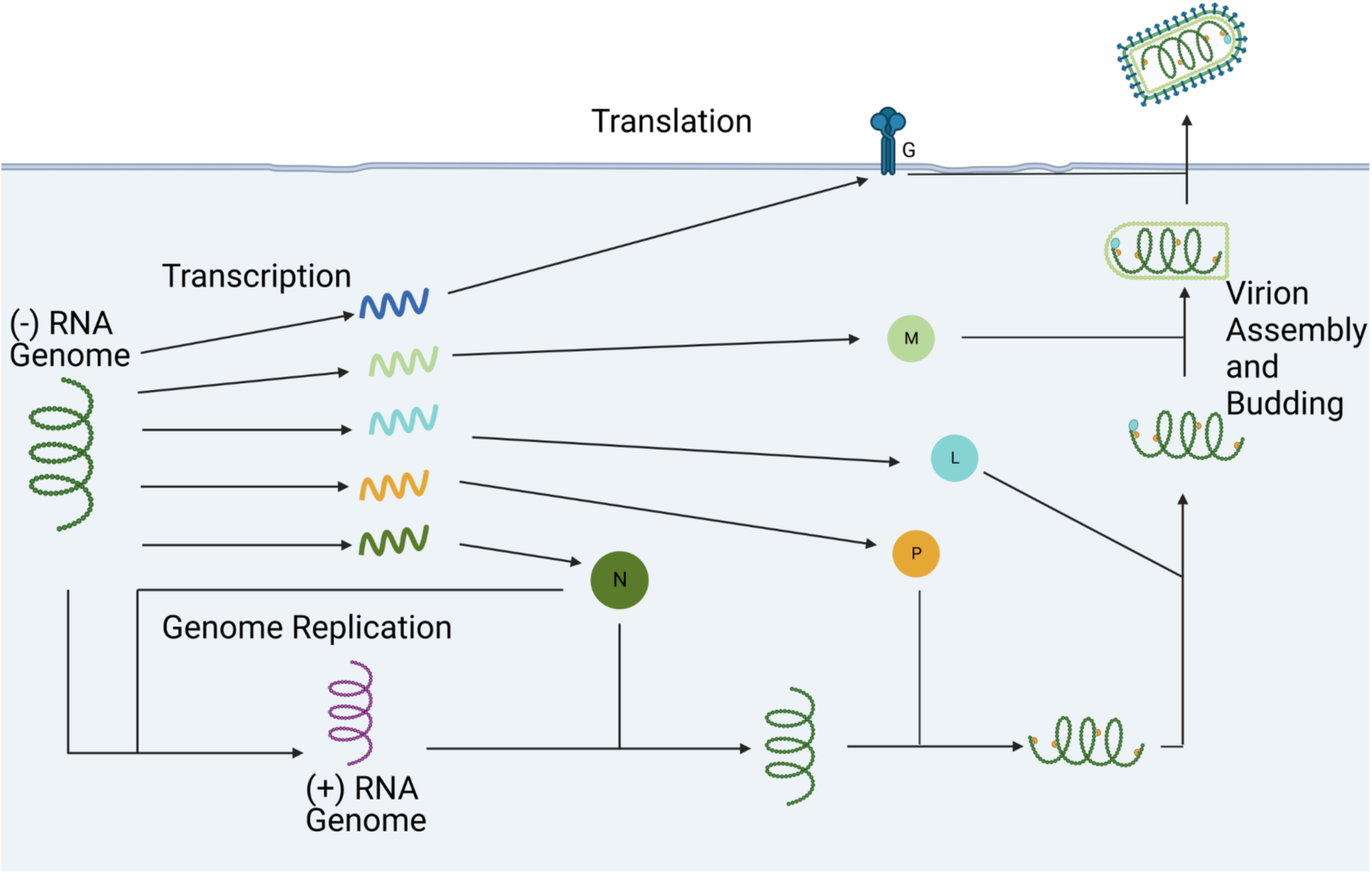
Simplified Model of Vesicular Stomatitis Virus. Visual representation of the simplified model used in this paper. Every molecule depicted has a degradation reaction associated with it that is not represented here.

### Transcription

The reactants in the transcription reactions are the nRNP species. This is based on the observation in the literature that the number of mN molecules produced is dependent on only the number of genomes present (12). In previous models, the transcription of VSV genes has been modeled so that they are coupled to the transcription of any upstream genes in the viral genome (7, 9). Here, we model the transcription reactions as independent from one another.

To calculate the transcription rates, we started by calculating the rate of transcribing the N gene. This was inspired from a previous stochastic model that split the transcription rate constants into the rate of polymerase recruitment and a delay to release the reactants (9). These were combined into a single rate constant. We then calculated the rate constants of all downstream genes by reducing the rate constant of each gene by 30% for each gene-gene junction upstream of the gene, except for the L polymerase which was reduced by 90% for the final gene-gene junction. This is to create the ratio of 1:0.7:0.49:0.343:0.0343 (ratio of mN to mRNA encoded in position 2 to mRNA encoded in position 3 to mRNA encoded in position 4 to mL) that is observed in the literature (23, 24).

### Translation

Each translation reaction is carried out independently and uses the corresponding mRNA to produce the protein. The rate constant of these reactions we use here assumes uniform saturation of the transcripts by ribosomes and is derived from a previous model (9). This uniformity of translation is also done to maintain the 1:0.7:0.49:0.343:0.0343 ratio, as it is also present at the protein level.

### Genome Synthesis

The reactants in the genome synthesis reactions are the pRNP species along with the N protein. This was calculated in a similar fashion to the transcription of the N gene by combining the estimated rate of polymerase recruitment with the delay to produce the antigenome (9). The decision to include the N protein in the reaction was because the N protein is known to play a pivotal role in facilitating and regulating genome/antigenome synthesis (10, 11).

### Antigenome Synthesis

The reactants in the antigenome synthesis reactions are the nRNP species along with the N protein. Since the amount of pRNP present in a cell is roughly 50 times higher than the amount of nRNP, we multiplied the genome synthesis rate constant by 50 as was done in a previous model (9).

### Degradation Reactions

All molecular species except for V have a degradation reaction associated with them. All rates are taken from ones used in a previous model (7).

### Model Simplification

Initially we attempted to simulate the binding of every protein to the genome as a separate reaction, but quickly realized that even when simulating the ODE model took an unreasonable amount of time. We then performed a series of tests where we increased the number of reactions and timed how the length of loading and simulating the ODE model changed as the number of reactions increased (S3 Dataset). We found that these times increased exponentially as we increased the reaction number (S1 Fig). This informed our ultimate decision that the best model to use is the simplest one. In this model, each molecule of protein represents a unit of the protein that makes up a virion. So, we then adjusted the rate constant of any reactions that involve the creation or consumption of proteins by dividing the rate constant by the number of proteins that is in a virion. For example, if the binding rate of protein P to the nsg is 0.0461 then it would be adjusted to be 0.0461/466 because 466 P proteins are in a single virion. A visual representation of this model, not including the degradation reactions, can be seen in Fig 1.

### Model Fitting

We then wanted to see if the parameters of the model could be adjusted to fit the mean number of virions produced by VSV gene shuffled variants that have been previously recorded in literature (19). We chose the data from BHK-cl.13 cells because it was the dataset where the mean number of viruses produced by the wild-type was closest to the mean number of viruses recorded when infecting single cells with wild-type VSV (25). We started by manually adjusting the parameters to get the wild-type model to be close to the mean number of virions produced by the wild-type VSV. This model, however, failed to reproduce the mean number of virions produced by the gene-shuffled variants (Table 2). These models can be found in the S4 Dataset. We then fit an ODE version of the model since this is much faster than fitting the stochastic model. This ODE model was able to reproduce the ranking of the variants (Table 2) but again did not replicate the mean number of virions produced by the different gene-shuffled variants. We then transitioned to a stochastic model that we could ultimately get to replicate the mean number of virions produced (Table 2). These models can be found in the S6 Dataset. This stochastic model allowed us to visualize the distribution in the number of virions produced by the wild-type model and see that it is consistent with viral distributions observed in the literature (S2 Fig) (25, 26, 27).

**Table 2.** Model Predictions and Mean Square Error (MSE).

| VSV Variant | Data From Literature | Manually Adjusted Param. | ODE-Model | Stochastic Model |
| --- | --- | --- | --- | --- |
| NMGPL | $4.35 * 10^3$ | $7.69 * 10^2$ | $4.19 * 10^3$ | $4.27 * 10^3 \pm 46$ |
| NPMGL (WT) | $2.09 * 10^3$ | $1.89 * 10^3$ | $2.48 * 10^3$ | $1.87 * 10^3 \pm 20$ |
| NMPGL | $2.35 * 10^3$ | $5.72 * 10^2$ | $2.69 * 10^3$ | $2.55 * 10^3 \pm 26$ |
| NGPML | $1.30 * 10^3$ | $3.05 * 10^3$ | $3.13 * 10^3$ | $1.33 * 10^3 \pm 10$ |
| NGMPL | $3.96 * 10^3$ | $1.90 * 10^3$ | $3.03 * 10^3$ | $3.93 * 10^3 \pm 41$ |
| MSE | | $4.66 * 10^6$ | $3.64 * 10^5$ | $1.92 * 10^4$ |

### Individual infections that exit the early phase of infection with more mRNA produce more virions

The individual trajectories of simulations using this model (Fig 2A) reveal heterogeneity in both the number of virions produced and the time at which the virions begin to be produced. Early on in infection, the random chance a reaction may or may not occur can have long lasting effects on the ultimate outcome of that infection, so we wanted to see if we could identify which early events contribute to the variation in the number of virions produced. We first set a distinction between the early phase of infection and the rest of the infection cycle by identifying when the first virion is produced. Here we define the time until the production of the first virion as the time it takes for the infection to leave the early phase. When we plot the time until the early phase is left against the number of virions produced (Fig 2B-F) we can see that there is a visible trend that the longer it takes for the early phase to end, the fewer virions that are ultimately produced.

**Fig 2.**
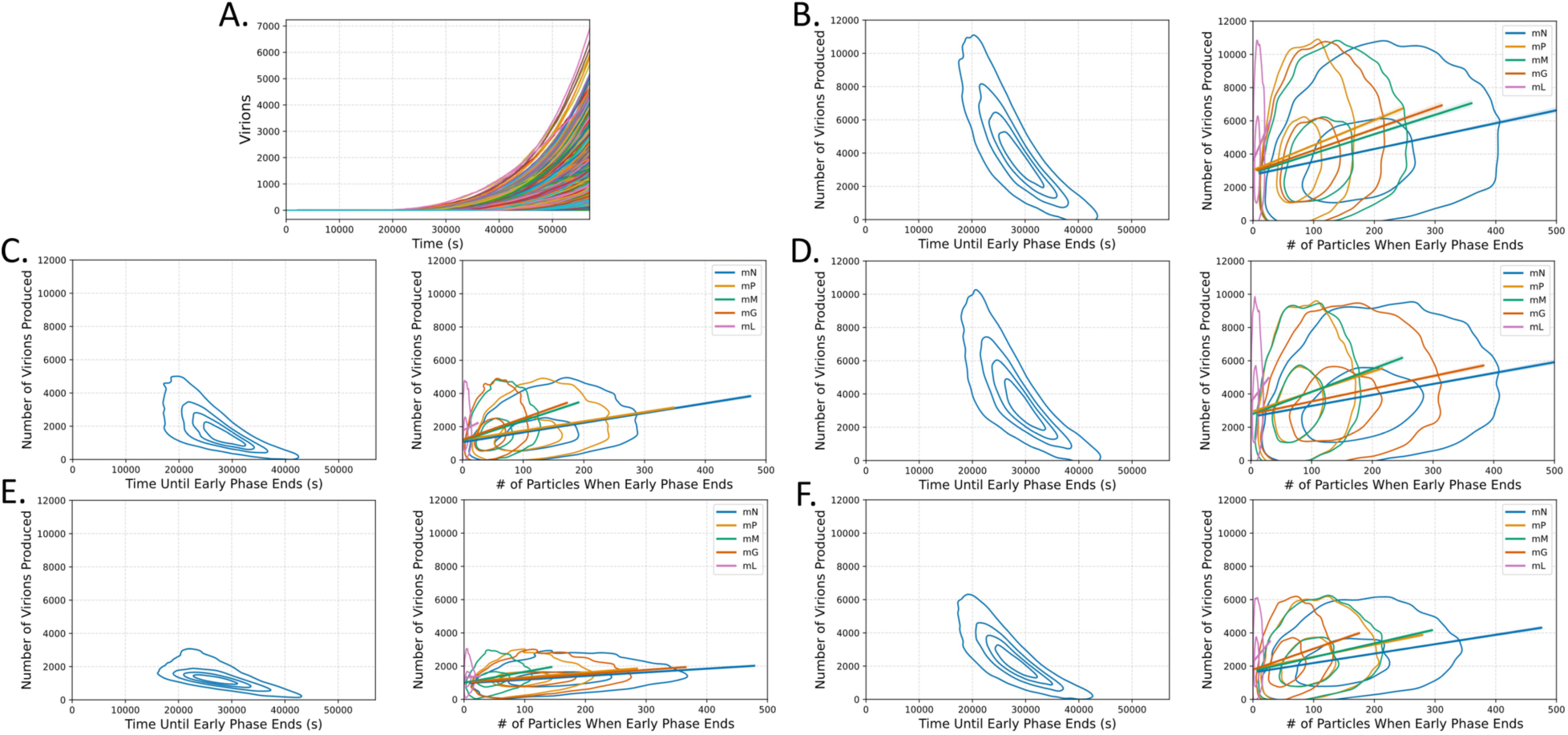
VSV infections that exit the early phase of infection earlier with more mRNA present produce more virions. (A) 1000 stochastic simulations of wild-type VSV showing the number of virions produced over time. (B-F) Left: Kernel density estimate (KDE) plots of the number of virions produced vs. the time until the early phase ends. Right: KDE plots of the quantity of each mRNA species present when the early phase ends, along with the 99% CI line of best fit, for variants (B) NMGPL, (C) NGMPL, (D) NMPGL, (E) NPMGL (wild-type), and (F) NGPML. All mRNA species show a weakly positive correlation (0.2 < r < .39) with the number of virions produced, except for mL which has a very weak correlation (0 < r < .19) in all variants.

This naturally then led us to investigate what early molecular species could also be participating in these observations. Naturally, we turned our attention to the quantity of mRNA present when the infection leaves the early phase, as the transcription reactions are some of the earliest reactions that occur upon infection. We plotted the number of mRNA present at the time when the infection exits the early phase against the number of virions produced (Fig 2B-F) and revealed a clear positive correlation between every mRNA and the number of virions produced. These observations together suggest that the variation we see in the number of virions produced within variants is likely due to variation in how long it takes for the infections to stockpile resources and actively start producing viruses. The script used to perform these analyses and generate the figures can be found in the S7 Dataset.

### The positions of the N, P, and L genes are the most important for maximizing virion production

We next turned our focus to identifying what is influencing the variation in the mean number of virions produced between different variants. We started by generating all 120 possible gene-shuffled VSV variants to observe the fitness landscape of these variants (Fig 3A). It appears that most gene-shuffled variants (102) are at least weakly viable (produce more than 1 virion per infection), but few produce virions on the scale of the wild-type virus (>1000). Even more intriguing, however, is that there are nine that are predicted to produce more virions than the wild type (Table 3).

**Fig 3.**
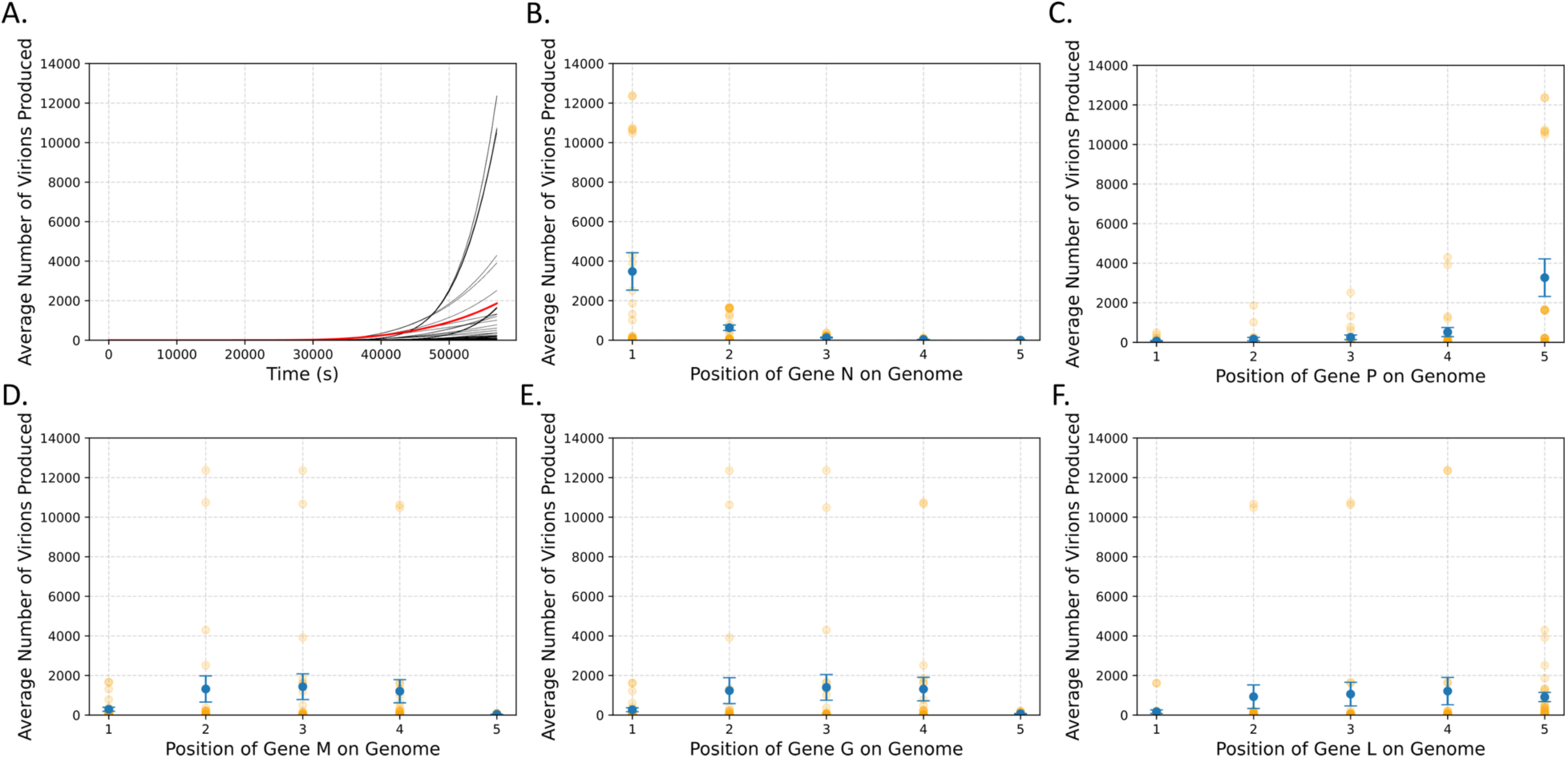
N and P gene position most strongly influence virion production in gene shuffled VSV variants. (A) The average number of virions produced over time for all 120 variants. The trajectory of the wild-type variant is in red. (B-F) The average number of virions produced as a function of gene position (which determines the transcription rate) for genes (B) N, (C) P, (D) M, (E) G, and (F) L. The error bars represent the standard error of the mean, and the orange dots represent the individual variants.

**Table 3.**
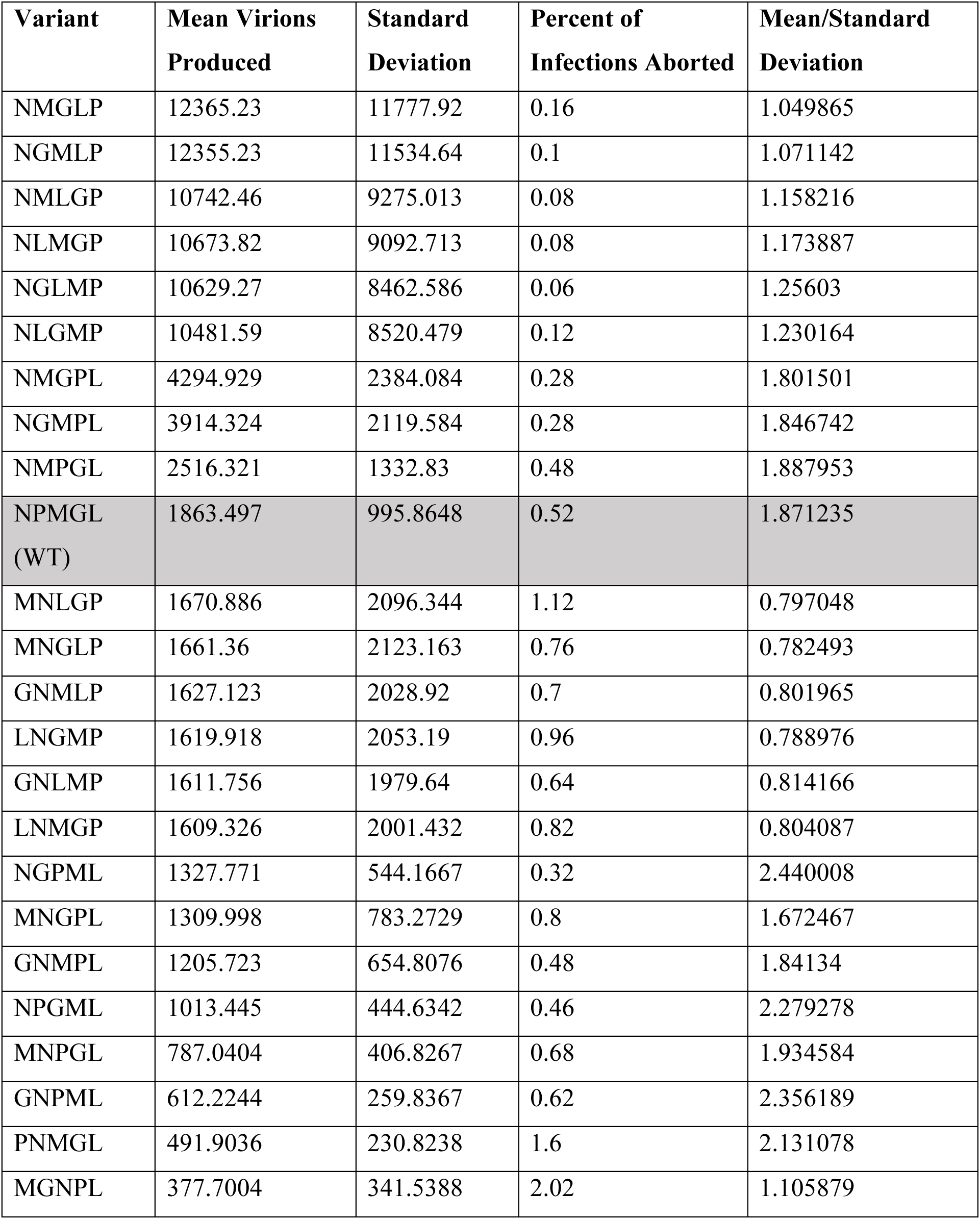
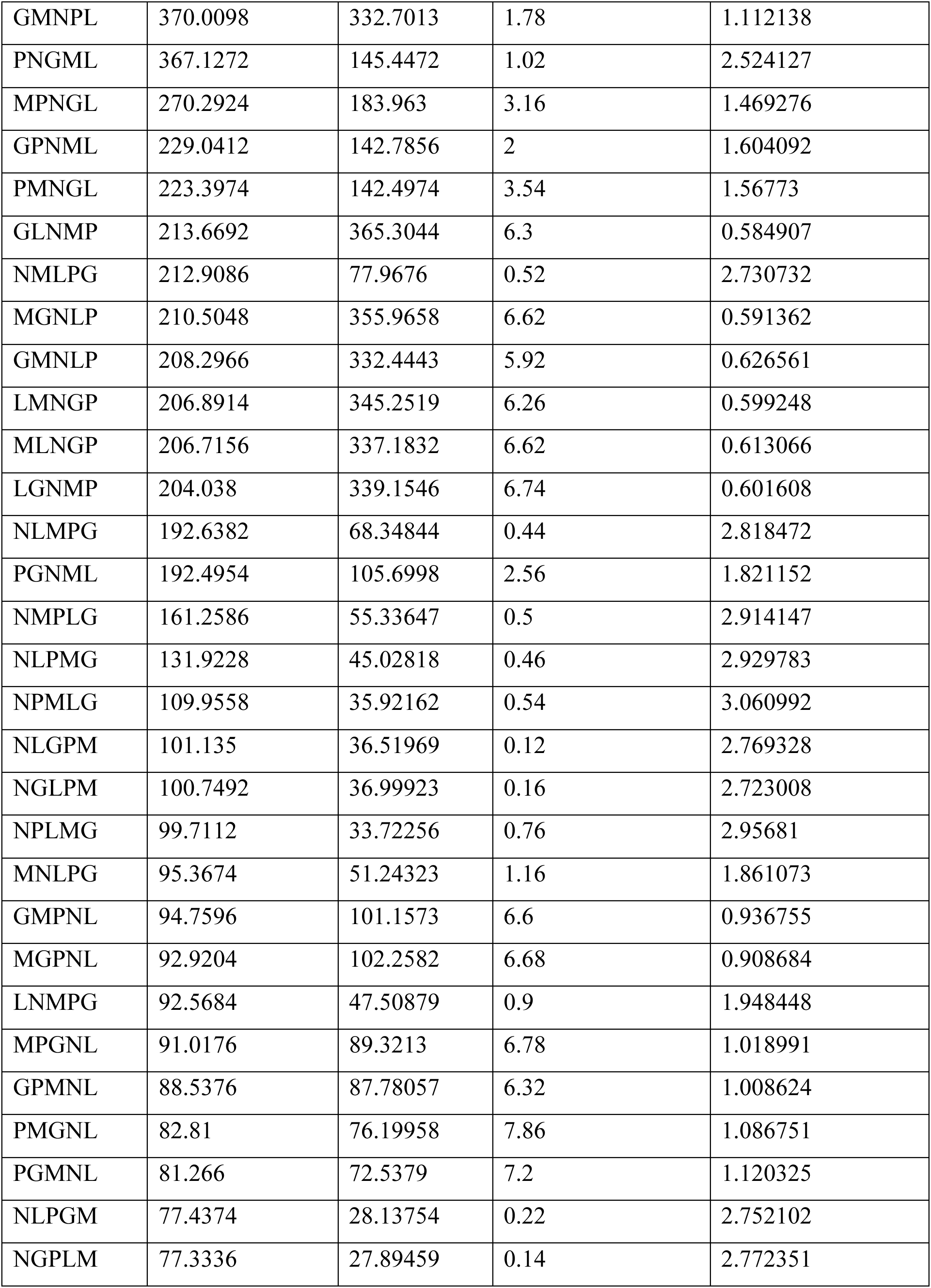
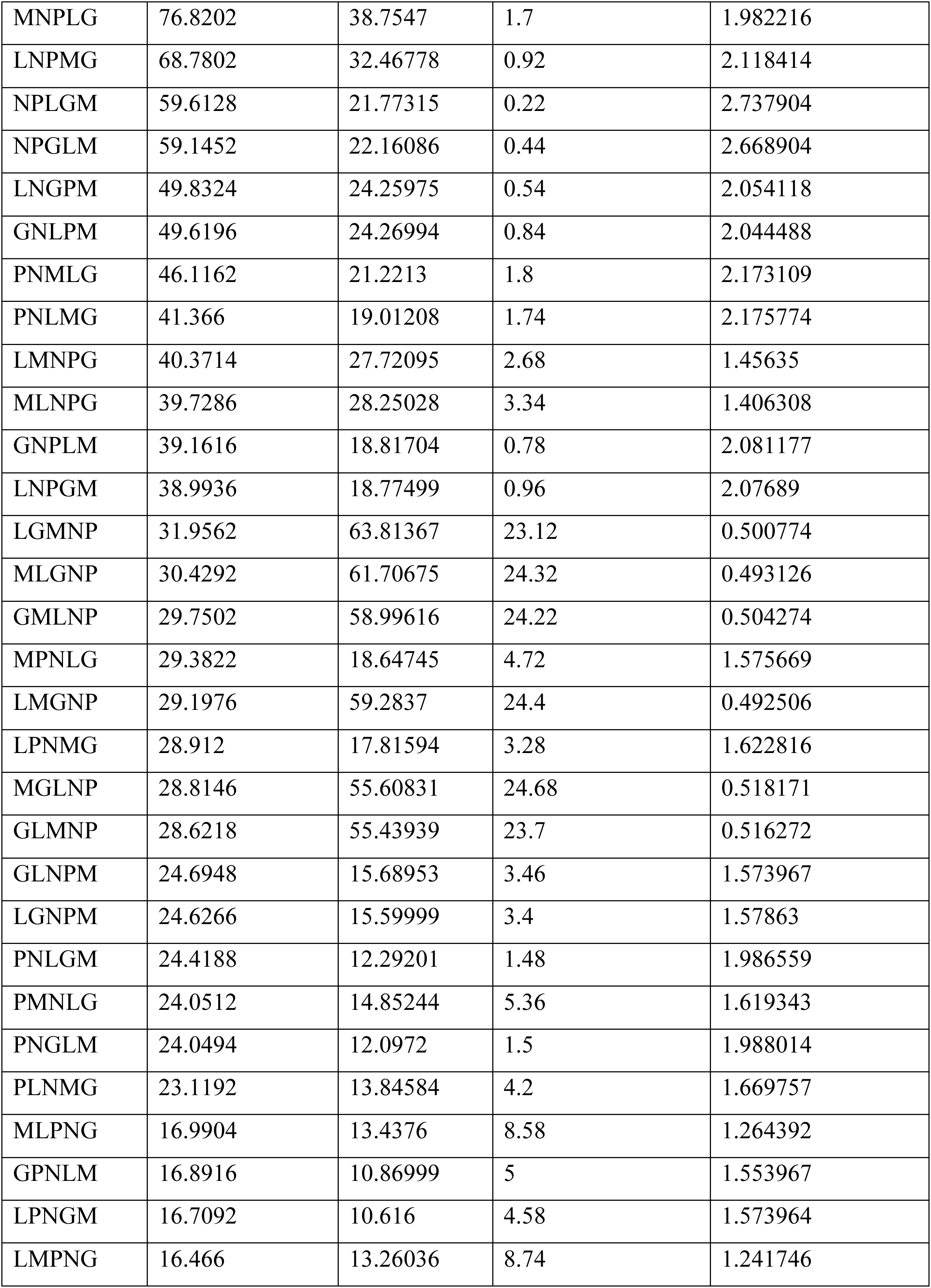
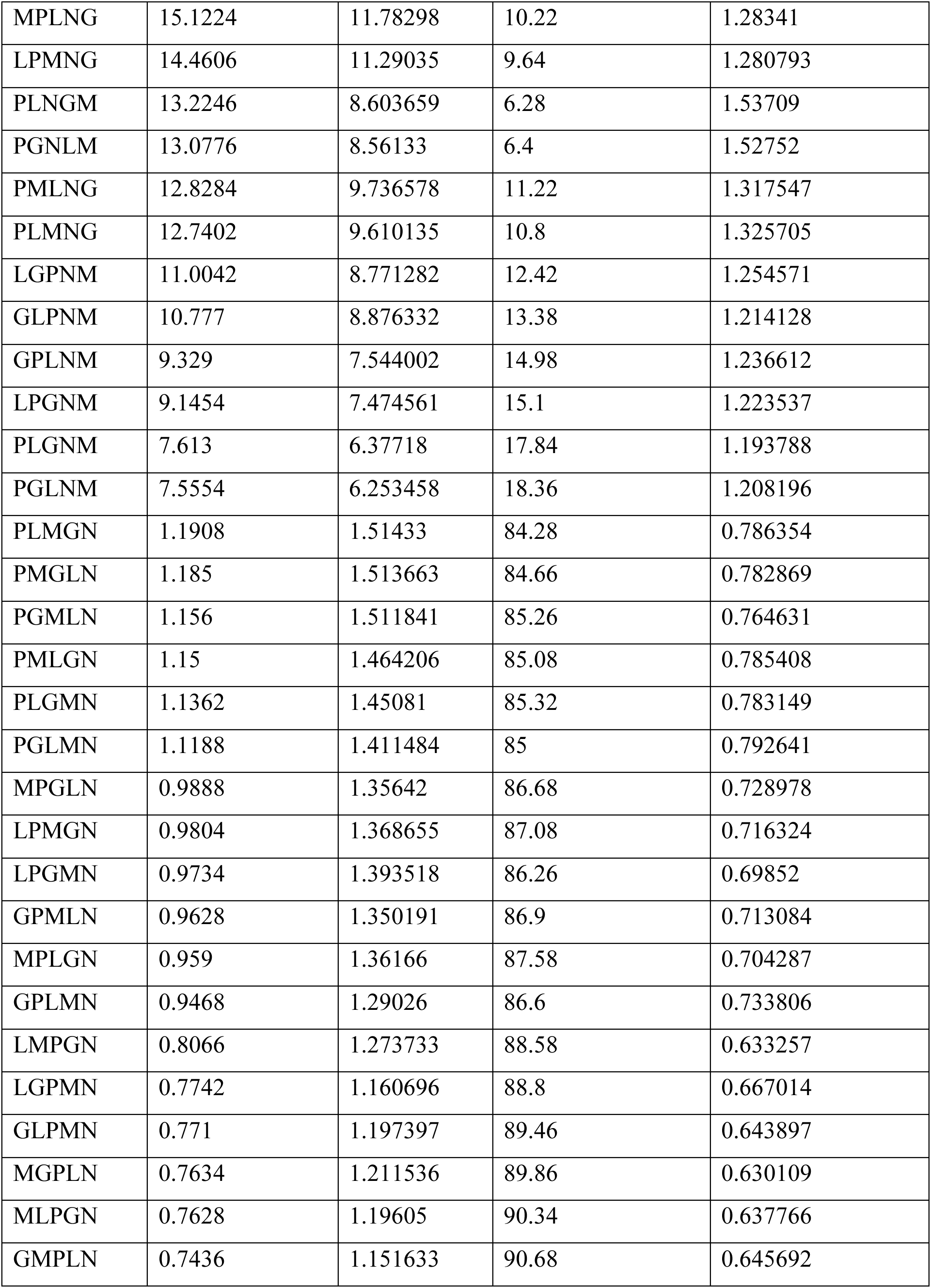
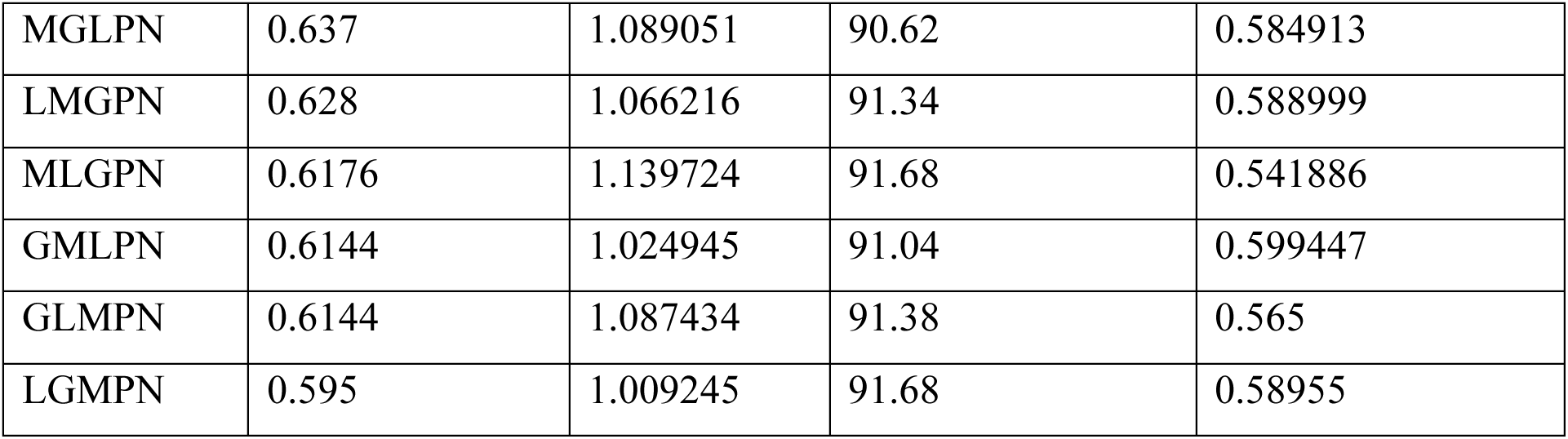
All Gene Shuffled Vesicular Stomatitis Virus Variants with Associated Statistics.

To identify what features made these variants so successful, we then looked at the average number of virions produced and the percent of infections aborted (infections producing <= 1 virion) by every variant as a function of each transcription rate to look for any patterns that may explain this variation (Fig 3B-F, Fig 4). For vitality, this revealed a negative correlation between the transcription rate of the N gene and the number of infections that were aborted. Most notably, when the N gene is in position 1, 2, or 3 nearly all infections are fruitful, while in position 5 nearly all infections are aborted. There appeared to be no obvious relation between the rate of aborted infections with the other genes, however (Fig 4).

**Fig 4.**
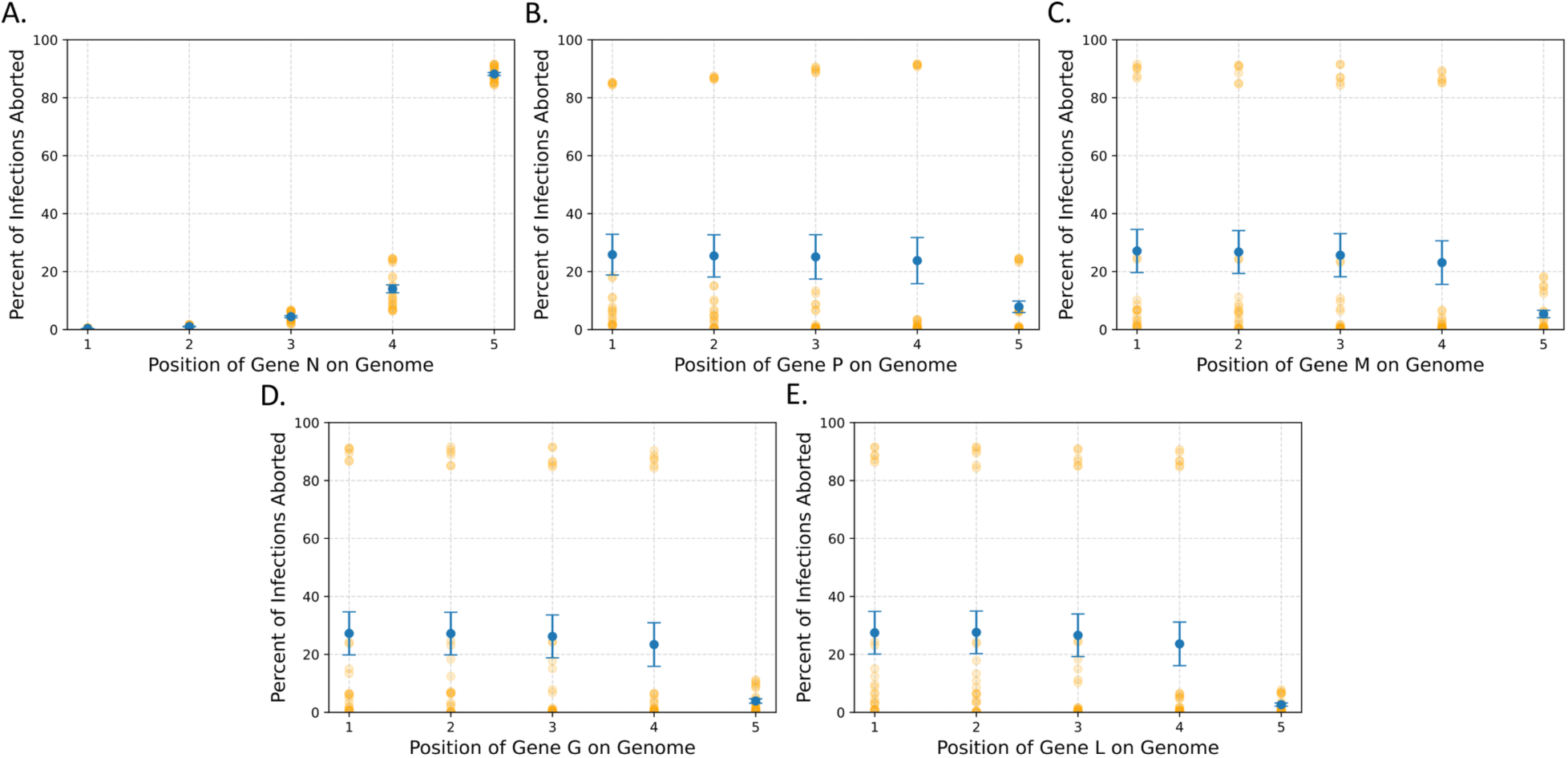
N gene position most strongly influences the percentage of infections that are aborted. The average percent of infections aborted as a function of the gene position of genes (A) N, (B) P, (C) M, (D) G, and (E) L for all 120 variants. The gene position is directly related to transcription rate of each gene. The error bars are the standard error of the mean and the orange dots represent the individual variants.

When looking at the number of virions produced, however, a much more complex behavior emerged. We observed a positive relationship with the transcription rate of N with the number of virions produced, but a negative correlation with the transcription rate of P. There also appeared to be a nonlinear relationship between the number of virions produced and the transcription rates of L, M, and G, all of which showed a small peak in the middle of the transcription rates. These observations are consistent with the gene orientation of the predicted most fruitful variant (NMGLP), as in this variant the N transcription rate is maximized, and the P transcription rate minimized with the other three in the middle.

These plots also revealed a few rules for what positions these different genes need to be in to produce a virus that yields more virions than the wild-type variant. The first is that the N gene must be in the first position of the genome. The next is the L and P genes must be in the 3^rd^, 4^th^, or 5^th^ positions, which is consistent with the idea that the transcription rate of P is negatively associated with the number of virions produced. Finally, the G and M genes must be either in the 2^nd^, 3^rd^, or 4^th^ positions, given the peak we observe for these middle rates.

These results suggest that the transcription rates of P and N appear to be the most influential on the number of virions each variant produce. This is further supported by the fact that all 6 variants that have N in the 1^st^ position and P in the 5^th^ position are the top 6 virion producing variants. However, it also appears that the position of L seems to be also important, as within these top 6 variants, if L is in the 4^th^ gene position the average yield is ∼12350 virions, while if it is in any other position, we only see ∼10000 virions produced (Table 3). In short, it looks like the best virion production occurs when the transcription of L and P are minimized and N is maximized. The script used to perform these analyses and generate the figures can be found in the S8 Dataset.

However, this only explains the variation in the most prolific phenotypes. It does not explain the wide variation in virion production of the other 14 variants that produce more than 1000 virions. This is because we are still left with only a fraction of an image of what the relationships between these transcription rates and the number of the virions produced are, because to alter one transcription rate in these models, you must alter at least one more.

### The balance of all transcription rates plays a role in determining virion production

The next step was to see how altering individual transcription rates independently could alter the number of virions produced. For this section we looked at the best variant and the wild-type variant to see how altering each transcription rate by values between −25% and +25% would affect virion production (Fig 5). The wild-type variant reveals that altering every transcription rate has potential to alter the number of virions produced. Here, it appears that the transcription rates of the genes encoding mN, mM, and mG have a positive association with the while the rates for genes encoding mP and mL have a negative association. The most fruitful variant (NMGLP), however, is only strongly altered by changing the transcription rate of mN which also has a positive association, but interestingly, has a very slight positive association with increasing the transcription rate of mP as well. The script used to perform these analyses can be found in the S8 Dataset.

**Fig 5.**
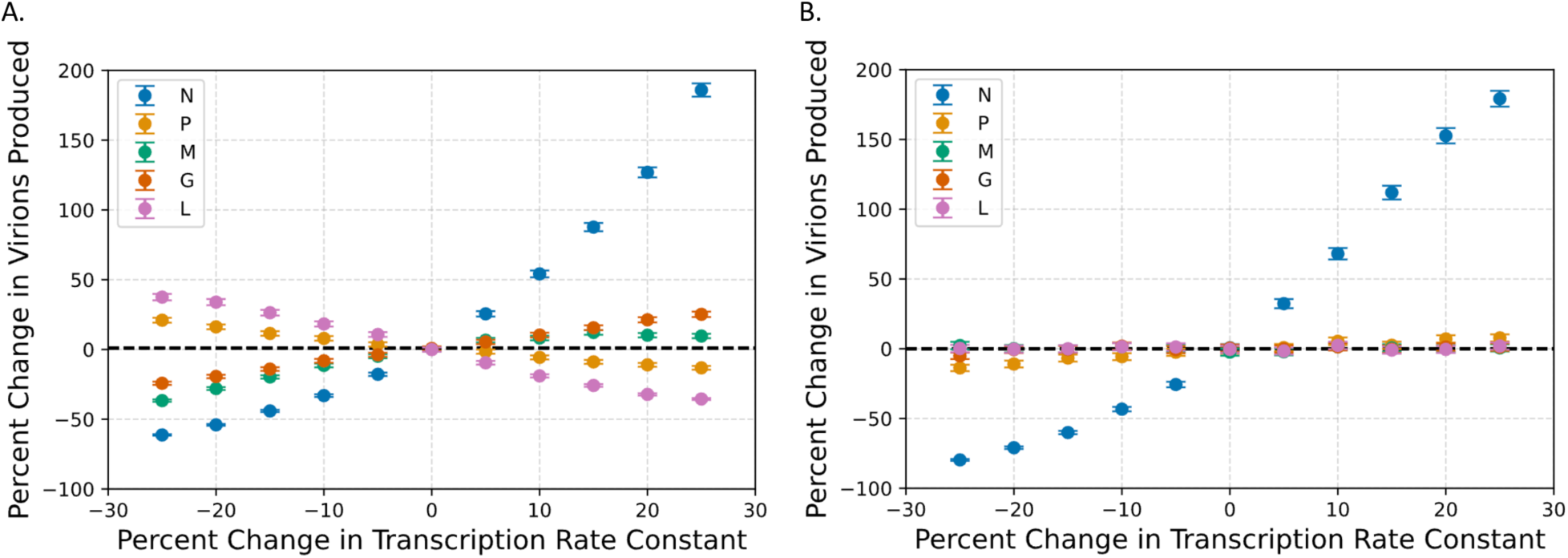
Parameter scans reveal the influence of L, M, and G genes on virion production. These graphs show how adjusting the transcription rates of each gene by a percentage of the value of the parameter in (A) WT and (B) the optimal variant (NMGLP) influences the number of virions produced. The black dotted line shows the mean number of virions produced by those variants.

These two observations together suggest that for each variant, there is an optimal balance of these transcription rates that maximizes virion production that is not simply achieved by maximizing all transcription rates. It appears that there is a hierarchy where the best virion production is achieved by maximizing N transcription but also by balancing the production of G, M, L, and P transcription, and deviations from this balance can alter virion production. This suggests that producing more virions is not achieved by merely maximizing transcription rates, but also by considering the interplay between all species.

## Discussion

In this study we sought to understand how the stochastic nature of the reactions in early cellular infections influence the variation we observe in the number of VSV virions produced by these cells. Previous models have attempted to investigate the cellular dynamics of VSV, but these models are either ODE-based (7, 12) or do not represent the full replication cycle due to the computational burden required to investigate the model (9). The model we present here is the first stochastic mechanistic model that captures the full replicative cycle of VSV. The model is also the most accurate one to date in predicting changes in virion production in response to gene shuffling.

This model provides valuable insight into the sources of variation in virion production, both within a single variant as well as between variants. Within variants, the main determinant of the number of virions produced appears to be how long it takes to leave the early phase of infection (Fig 2B-F). This is consistent with the observation that some of the variability in virion production can be explained by what phase in the cell cycle the cell is infected in (25). VSV’s mechanism for killing cells is tied to the cell cycle, and so the phase the cell is in could influence the ultimate number of virions that can be produced by determining how long VSV has to start replicating (28). Here, the variation in the amplifying time of VSV is the result of delays in entering the replicative cycle, which may allow us to capture some of the variation that is due to infecting cells at different cell cycle stages. Nonetheless, this intrinsic feature we identified is likely to play a role in intercellular variation. What is also exciting is that we can capture these delays by implementing the traditional stochastic simulation algorithm without the implementation of a DSSA.

As for the variation between VSV variants, our model seems to suggest that the most important factor for determining the number of virions produced is the transcription rate of the N gene. This is evident as all variants that produce more than 2000 virions have the N gene in the first position (Fig 3B). In addition, when we increased the transcription rate of each gene independently, N led to the largest increase in the number of virions produced by both the wild-type and most fruitful gene shuffled variant (NMGLP) (Fig 5). Furthermore, the transcription rate of the N gene appears also to play a major role in how frequently infections are aborted (Fig 4). These findings are consistent with observations in the literature that genome and antigenome replication are both strongly dependent on and regulated by the N protein (10, 11). As we increase the transcription rate of the N gene, we likely also increase the rate of replication in the cell, producing more genomes and ultimately, exponentially more virions.

However, we also see that the other genes also actively participate in determining the virion output, although to a lesser extent. There is clearly a complex balance of transcription rates between M, G, L, and P that allows for the optimal growth rate, and deviations from this optimal balance can alter virion production. This is likely due to the distinction between active vs. inactive nsg in the model. Since L and P both bind to the active genome, excess L and P might cause newly synthesized genomes to be converted into the inactive form too quickly. This may explain the negative correlation between these transcription rates and virion production in the wild-type. Indeed, there is evidence that the P protein may inhibit transcription (29). M and G then convert those inactive genomes into virions, and as such, they can influence virion production by forming bottlenecks during virion assembly. L and P have the potential to do this as well, and so it may be that maximizing virion production is a matter of balancing how long nsgs stay in the active state with how quickly they are incorporated into virions.

The negative relationship of the transcription rate of the P and L genes and the positive relationship of the transcription rate of the N gene with regards to virion production are consistent with the results of a previous model that also sought to predict how all 120 variants of VSV may behave (8). However, our predictions appear to put a heavier weight on the transcription rate of P, while the previous model put more emphasis on L. As for the M and G genes, both our and their results support the notion that maximum virion output is achieved when these are in gene positions 2, 3, or 4, and that when M and G are in position 5, almost no virions are produced. However, our results differ in that putting these genes in position 1 also is more detrimental than they predicted and is similar to putting them in position 5.

As for the overall fitness landscape of the variants, they also predicted that several variants produce as many virions as the wild-type as well as a few that produce more than the wild-type (8). However, the different relationships between transcription rates we found alter which variants are in these positions. For example, their predicted best gene shuffled variants are the NMPGL and NMGPL variants, while our predicted best variant is the NMGLP variant. Likely these differences are due to differences in the datasets utilized for fitting the model, as we fit the mean number of virions produced to the mean number of virions produced per cell in the five VSV gene shuffled variants, while their model was fit to ranked variant data of six gene shuffled VSV variants from a previous paper (30) and their own one-step growth curve of the wild-type. Furthermore, the cell type of the ranked variant data was from BSC-1 cells, while the data we utilized was from BHK-cl.13 cells. This is extremely important as the number of virions produced and the ranking of variants is highly dependent on the cell line utilized to grow the virus (19). For example, in our dataset of the five gene-swapped variants, the wild-type was the fourth highest producing variant, while in the dataset they used, the wild-type was the third highest producing variant of six gene shuffled variants.

However, since our model is stochastic and the previous model is an ODE model, our model has additional traits that we can observe in the variants that are not possible in the ODE model. We can observe the distribution of the number of virions produced by different variants (S2 Fig) as well as predict how frequently infections may be aborted (Figure 4). We can also calculate additional statistics about the virion distribution, allowing us to select for more complex phenotypes when designing specific viral behaviors. For example, if we wanted to maximize the mean number of virions produced while minimizing the standard deviation, we could calculate the ratio of mean to standard deviation (Table 3). Our model currently predicts that the best variant for achieving such a phenotype would be variant NPMLG. These attributes of our model allow for more in-depth analyses of the effects of different mutations on the viral life cycle than any previous model beforehand.

## Methods

Model creation was formalized as previously described (31).

### Model Creation

The models were encoded in XML format. These files were generated by creating two Excel spreadsheets, one containing the reactions and their parameters and the other containing the initial number of molecular species. The models were then fed to the Python script provided in S1 Dataset.

### Model Simulation

Models were simulated using COPASI’s Python implementation, COPASI Basico (32). ODE models were simulated using the LSODA Method, while stochastic simulations were performed uing the Gillespie Direct Method. Models were simulated for ∼16 hours (57,000 seconds). This time was chosen because there is not a solid consensus on how long VSV takes to kill a cell, but it is known that virions start emerging around 2-6 hours after infection (33) and that VSV’s mechanism for killing cells is tied to the cell cycle (28). So, if we assume the average cell cycle time in the cells we are modeling (34) is about a day and that the cells are asynchronous, then an average time until death of 16 hours should be reasonable. The dataset used to analyze the stochastic models consisted of 10,000 trajectories of each variant.

### Parameter Optimization

Models were fit using the minimize function from SciPy (35). The Nelder-mead method was utilized. ODE and stochastic Models were fit by calculating the mean squared error (MSE) of the number of virions present at the endpoint (57,000 s) and the average number of viruses produced recorded in the literature. The scripts used to fit the model can be found in S5 Dataset.

### Cloud Computing

When performing large numbers of optimizations or stochastic simulations, we utilized AWS’s EC2 service to create an instance that could handle the computations. We utilized a single c5.18xlarge instance.

